## supplement for "Emergence of alternative offspring care patterns from the evolution of parental negotiation strategies"

This Supplement includes:

**Fig. S1:** Evolution of parental provisioning levels in the baseline model

**Supplementary text:** Fitness considerations and mathematical analysis of the baseline model

**Supplementary references**

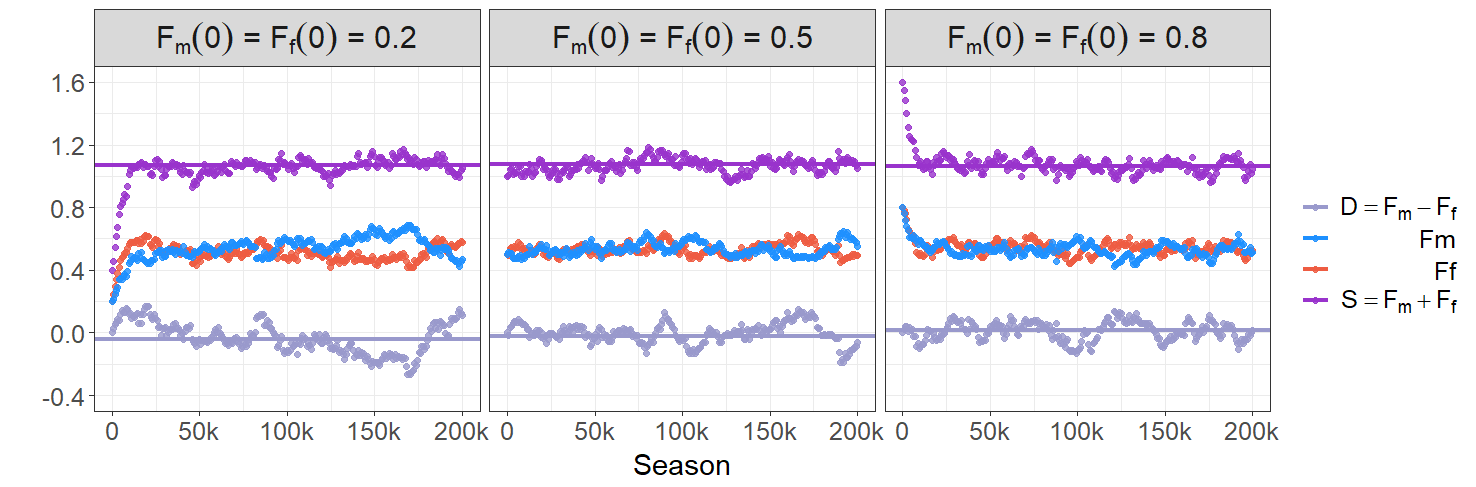

**Figure S1: Evolution of parental provisioning rates in the baseline model.** In the baseline model, the parental provisioning rates (
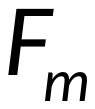
 and
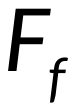
 for the male and female parent, respectively) do not result from negotiation but are heritable parameters that are fixed throughout an individual’s lifetime (see Methods). The three panels show three representative simulations of the evolution of
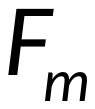
 (blue curves) and
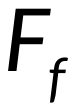
 (red curves) for three initial conditions. As in hundreds of other simulations,
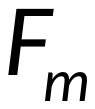
 and
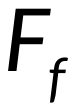
 rapidly converged to an equilibrium where for our default parameters both parents exhibit a provisioning rate of about
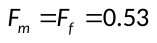
 feeds per minute, resulting in a total provisioning rate of about
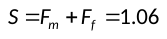
 (purple curves). While
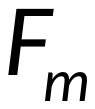
 and
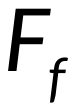
 fluctuate around their equilibrium value in a complementary manner, the total provisioning rate *S* remains relatively constant. The grey curves show the difference in parental provisioning rates,
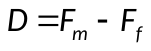
, which fluctuates around zero. All simulations run for the baseline model resulted in egalitarian biparental care with approximately the same total provisioning level.

**Supplementary text: Fitness considerations and mathematical analysis of the baseline model**

***The Houston-Davies model***

Our baseline model may be viewed as an individual-based implementation of the classical model of Houston and Davies (1985), a foundational contribution to parental care research. As their point of departure, Houston and Davies considered a fitness function of the form

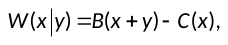
(S1)

where *x* is the parental effort of a focal parent, *y* is the parental effort of its partner,
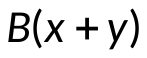
 is the fitness benefit accrued via the current brood, and
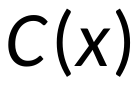
 is the fitness cost associated with the effort of the focal parent. Functions B and C are assumed to be increasing functions, with some additional assumptions about their shape. The fitness function is a prototype illustration of sexual conflict over parental care, as the fitness benefits increase with the efforts of both parents, while the costs only depend on a parent’s own effort. Accordingly, both parents have an (evolutionary) interest in letting the partner do most of the work. Given the fitness function (S1), Houston and Davies (and others, e.g. McNamara et al. 1999; Johnstone et al. 2014) derived conditions to be satisfied by evolutionarily stable parental strategies and, from these, drew conclusions on the properties of these strategies.

The Houston-Davies model was a major conceptual advance, which inspired a lot of follow-up work. However, in concrete applications, the determination of the ingredients of the fitness function is often not straightforward. The two components are often interpreted in terms of reproductive values:
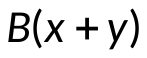
 corresponds to the reproductive value of the current brood, while
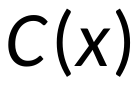
 is related to the residual reproductive value of a parent employing effort *x*. While there are standard methods to calculate reproductive values for several classes of models (e.g., Houston & McNamara 1999; Caswell 2001; Otto & Day 2007), these models are often “ecologically inconsistent” in that they do not include population regulation (and, hence, assume exponentially increasing or declining populations). Including population regulation in the model is important, as the mechanism of population regulation can have major implications for the course and outcome of evolution (e.g., Mylius & Diekmann 1995; Pen & Weissing 2000). In many examples, it may be straightforward to include population regulation in the Houston-Davies model, but even in these cases, the resulting fitness function is more complicated than the simple form (S1). A second complication arises because most parental care models consider the interaction between two sexes. This necessitates distinguishing between male and female reproductive values and the inclusion of the consistency requirement (the “Fisher condition”) that the reproductive output of all males in the population exactly matches the reproductive output of all females (Houston & McNamara 2002; Wade & Shuster 2002; Fromhage & Jennions 2016). Although this can be done (e.g., Pen & Weissing 2002), the resulting fitness function is more complicated than equation (S1).

The take-home message of this is that fitness functions of the Houston-Davies type have led to considerable conceptual clarification, but that the fitness functions of ecologically consistent two-sex models of parental care should not be expected to have the simple form (S1). As a consequence, the conclusions drawn from the Houston-Davies model cannot immediately be extrapolated to real-world scenarios.

***Individual-based evolutionary models***

In individual-based models, the Fisher condition is automatically satisfied. Moreover, such models must be designed to be ecologically consistent; otherwise, the modelled populations would explode. In comparison to the more standard mathematical models, individual-based evolutionary simulation models have the advantage that evolution through natural selection can be studied without specifying a fitness function (which, as in the Houston-Davies model, is often the starting point of mathematical models). To be sure, assumptions have to be made on “fitness components” like survival probabilities or fecundities, but these do not have to be integrated into a unified measure of “fitness”.

Individual-based models have the additional advantage that the evolutionary dynamics is built into the model. To be sure, this dynamics is affected by model assumptions regarding population regulation (see above), the population size (affecting genetic drift), the mutation process (affecting diversity and the mutation-selection balance), the genetic architecture underlying the heritable strategies (affecting genetic recombination and the co-evolution of strategies), and the reproductive system (e.g., sexual versus asexual). However, these assumptions are typically more transparent than the more implicit assumptions underlying mathematical models. For example, mathematical models often assume that evolution through natural selection proceeds in the direction of the “fitness gradient”, that is, in the direction of the steepest ascent of the fitness function. As shown by Long and Weissing (2023), this assumption can lead to wrong conclusions in the context of parental care evolution.

***Mathematical analysis of the baseline model***

Relatively simple individual-based models, like our baseline model, are, to a certain extent, amenable to mathematical analysis. As shown in Long & Weissing (2023), such analysis is useful as it sheds light on the pros and cons of the various tools used to investigate natural selection. Therefore, we here sketch the fitness analysis of our baseline model. As a first step, we have to consider population regulation. In our model, the population size is regulated by the fact that juveniles can only enter the population of adults if positions become available through adult mortality. Hence, in our model, the juveniles (and not the adults) are affected by density dependence. As shown in Mylius & Diekmann (1995), this implies that “expected lifetime reproductive success” is an adequate fitness measure.

In our model, expected lifetime reproductive success (ELRS) is given by the product of two factors: the expected number of offspring produced in a reproductive season and the expected number of reproductive seasons. As explained in the Methods section of the main text (eqn (4)), the expected number of offspring produced per reproductive season is proportional to
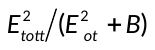
, where
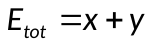
 is the total effort of both parents. The expected number of reproductive seasons of an individual employing effort *x* is proportional to the individual’s life expectancy, which in turn is given the inverse of the individual’s per-season mortality. As in our model the probability of a parent with effort *x* to survive the season is
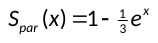
 (see eqn (3) in the Methods section of the main text), the life expectancy of an individual with effort *x* is given by:
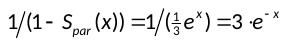
. Taking the product of the two factors, we conclude that the ELRS of an individual employing effort *x* in a population where the parental partners tend to employ effort *y* is proportional to:

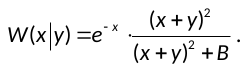
(S2)

While in the Houston-Davies model (S1) the reproductive costs are subtracted from the reproductive benefits, ELRS in our model corresponds to the quotient of a benefit term
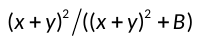
 (corresponding to per-season fledgling production) and a cost term
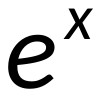
 (corresponding to per-season mortality). From (S2), the optimal parental effort can readily be determined. Inspection of the derivative of
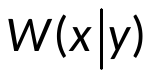
 with respect to *x* reveals that it has the same sign as the cubic

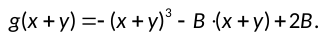
(S3)

Hence, the derivative of
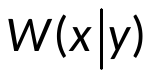
 with respect to *x* is zero if
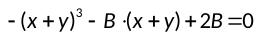
, which for our choice of *B* (
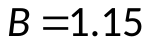
; see Methods) is solved by
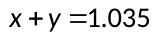
, or
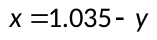
. As the cubic *g* is a declining function, it must be positive to the left and negative to the right of this value of *x*. Accordingly, for a given value of *y* the fitness function
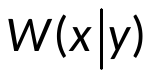
 increases for
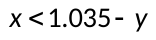
 and decreases for
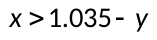
. In other words,
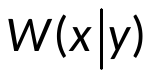
 is maximised for
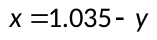
; this parental effort is the “best response” of an individual if its interaction partners employ effort *y*.

From this, we can conclude that natural selection has the tendency to achieve a total effort
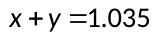
, a value that agrees reasonably well with the equilibrium total effort
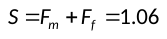
 we found in our simulations of the baseline model (see Fig. S1). The discrepancy between the two values may be explained by the fact that until now we did not consider any sex differences in the derivation of our fitness function (S2). In our model, the sexes do not differ in their life history characteristics, and the simulations in Fig. S1 reveal that parental care is pretty egalitarian. Yet, most of the time the difference
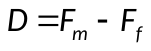
 of male and female parental efforts is not equal to zero, implying that one sex invests (slightly) less in the current brood than the other sex. This has two implications (Long et al. 2024). First, the more-caring sex has a (slightly) higher mortality, reducing its life expectancy and, hence, its expected lifetime reproductive success. Second, this higher mortality results in a bias in the adult sex ratio: the less-caring sex becomes overrepresented in the population. As a consequence, not all members of the less-caring sex can find a mating partner in a given season, implying that their longer life expectancy does not result in more matings and, hence, a higher lifetime reproductive success (see Long et al. 2024 for details).

All these ramification could be included in a sex-differentiated fitness function (as in Pen & Weissing 2002), but we refrain from deriving such a function in the current effort.
